# Selective breeding for determinacy and photoperiod sensitivity in common bean (*Phaseolus vulgaris* L.)

**DOI:** 10.1101/2024.10.27.620491

**Authors:** Kate E Denning-James, Caspar Chater, Andrés J Cortés, Matthew W Blair, Diana Peláez, Anthony Hall, Jose J De Vega

## Abstract

Common bean (*Phaseolus vulgaris* L.) is a legume pulse crop that provides significant dietary and ecosystem benefits globally. We investigated two key traits, determinacy and photoperiod sensitivity, that are integral to its management and crop production, and that were early selected during the domestication of both Mesoamerican and Andean gene pools. Still, significant variation exists among common bean landraces for these traits. Since landraces form the basis for trait introgression in pre-breeding, understanding these traits’ genetic underpinnings and relation with population structure is vital for guiding breeding and genetic studies.

We explored genetic admixture, principal component, and phylogenetic analyses to define subpopulations and gene pools, and genome-wide association mapping (GWAS) to identify marker-trait associations in a diversity panel of common bean landraces. We observed a clear correlation between these traits, gene pool and subpopulation structure. We found extensive admixture between the Andean and Mesoamerican gene pools in some regions. We identified 13 QTLs for determinacy and 10 QTLs for photoperiod sensitivity, and underlying causative genes. Most QTLs appear to be firstly described. Our study identified known and novel causative genes and a high proportion of pleiotropic effects for these traits in common bean, and likely translatable to other legume species.

**Highlight:** We identified and explored QTLs for the domestication-related determinacy and photoperiod sensitivity traits, which are traits critically associated with population structure and management and crop production.

## Introduction

Common bean is a global staple that provides significant dietary and economic services by improving health and nutrition whilst helping to reduce poverty, specifically in developing countries. Common beans have also been labelled as one of the essential crops to mediate climate change due to their lower environmental impact and protection of food and nutritional security [1]. Common beans are cultivated mainly as grain legumes, but the immature seeds, pods and leaves are also eaten [2, 3]. There are hundreds of varieties, and the prevailing type grown in a country depends on market preferences [4]. Common beans are rich in essential dietary components, such as protein, minerals, fibre and micronutrients [3, 5–7], and protect against some forms of malnutrition, including stunting in children and micronutrient deficiencies [3, 8–10]. As legumes, common beans have a symbiotic relationship with nitrogen-fixing bacteria, allowing them to fix atmospheric nitrogen and enhance nitrogen levels in the soil, thereby reducing the need for expensive chemical fertilisers whilst improving yields [11–14]. Despite its widespread usability, trait segregation within and among bean landraces is still widespread, especially for critical agronomic traits such as growth habit and photoperiod.

Common bean underwent two separate domestications resulting in two gene pools: Andean and Mesoamerican. In addition, there are different races, intermediate species and admixed accessions due to genetic isolation, fragmentation and artificial selection for different morphological traits. The gene pools of common beans grow in a large variety of environments in the neotropics. This ecogeographic conditions, together with isolation by distance, have disrupted the gene flow between wild and domesticated common beans, and between the different gene pools [15, 16]. Consequently, there are large differences in their life history traits, morphology and genetics [16–19]. Another difference is cultivars are commonly autogamous and annual, while wild common beans and related species can be perennial and allogamous [20–22].

Photoperiod insensitivity and determinacy arose separately in both gene pools during the domestication of common beans, likely co-selected by growers [23, 24]. Wild common beans tend to be indeterminate and photoperiod-sensitive, requiring a particular day length to flower. Indeterminate growth is advantageous in the wild due to competition with surrounding vegetation, while photoperiod sensitivity was likely reinforced by divergent natural selection and local adaptation. On the other hand, photoperiod insensitivity was selected (likely unconsciously) as cultivated common beans were spread along a greater range of latitudes and environments. Determinacy, a developmental feature that causes common beans to have a terminal inflorescence when switching to a reproductive state [25], optimised agricultural management and harvesting efficiency. Determinate common beans tend to have a bush growth habit with reduced branching and vining abilities compared to the indeterminate varieties [26], therefore translocating biomass resources into an increased fitness output. While indeterminate and photoperiod sensitive landraces are common, the combined selection for photoperiod insensitivity and determinacy resulted in common bean varieties with shorter flowering periods, earlier maturation and easier management during harvesting [27, 28]. Photoperiod insensitivity and determinacy are advantageous traits from an agronomical point of view due to earlier harvesting and shorter exposure to unfavourable weather patterns under climate change, consequently providing better food security for communities [29, 30].

Modern breeding programmes are moving beyond a yield-centred paradigm to target resistance to biotic and abiotic stress, and also nutritional quality [31–34]. Landraces and crop wild relatives offer a promising reservoir of genetic diversity for these traits by introgression from the landraces into the elite genetic background [35–38]. However, understanding the genetic diversity, population structure, patterns of adaptations, and how these correlate with determinacy and photoperiod insensitivity is required to guarantee the retention of these key domesticated traits within future breeding cycles, given their association with crop management and production [16].

Common beans in Colombia are diverse regarding growth habits and photoperiod sensitivity. Colombia is the northernmost part of the Andean gene pool and south of the Mesoamerican and may act as a region of confluence between them. Consequently, it has been proposed that the region has a large amount of admixture and introgressive hybridisation [39–42]. Admixture and hybridisation lead to introgressions from differential parental origins, introducing new alleles and novel epistatic interaction into a population, allowing for new trait combinations that could merge exotic variation from diverse germplasm with more agronomic-desirable traits such as determinacy and photoperiod insensitivity.

Considering the above hypothesis, we characterised 144 representative landraces from Colombian and neighbouring countries, together with controls from other regions, using whole-genome re-sequencing. We utilised genome-wide association mapping (GWAS) to identify significant SNPs for photoperiod insensitivity and determinacy in this diversity panel. The novelty of this work lies in that prior research commonly focused on the Mesoamerican diversity rather than the Andean, due to the greater genetic diversity in the former, and had ignored admixed materials as an essential source of variation. Furthermore, research has rarely utilised whole genome sequencing of common bean accessions to undertake a GWAS on determinacy and photoperiod insensitivity phenotypes. Instead, previous work has mostly used QTL mapping and low density marker panels, resulting in poor resolution [28, 43, 44].

## Materials and Methods

### Diversity panel

The diversity panel was comprised of 144 genotypes mainly from Colombia and surrounding countries in Central and South America (Figure 1). The panel contained accessions from elite backgrounds, landraces, heirlooms, weedy and wild materials. The material was sourced from the International Centre for Tropical Agriculture (CIAT)’s genebank, the Leibniz Institute of Plant Genetics and Crop Plant Research (IPK)’s genebank, and heirlooms bought from the catalogues from ‘Jungle Seeds’ [45] and ‘Beans and Herbs’ [46] in 2020. The panel represents the Andean and Mesoamerican gene pools and races, as well as diverse seed coat colours and Colombian varieties.

**Figure 1:**
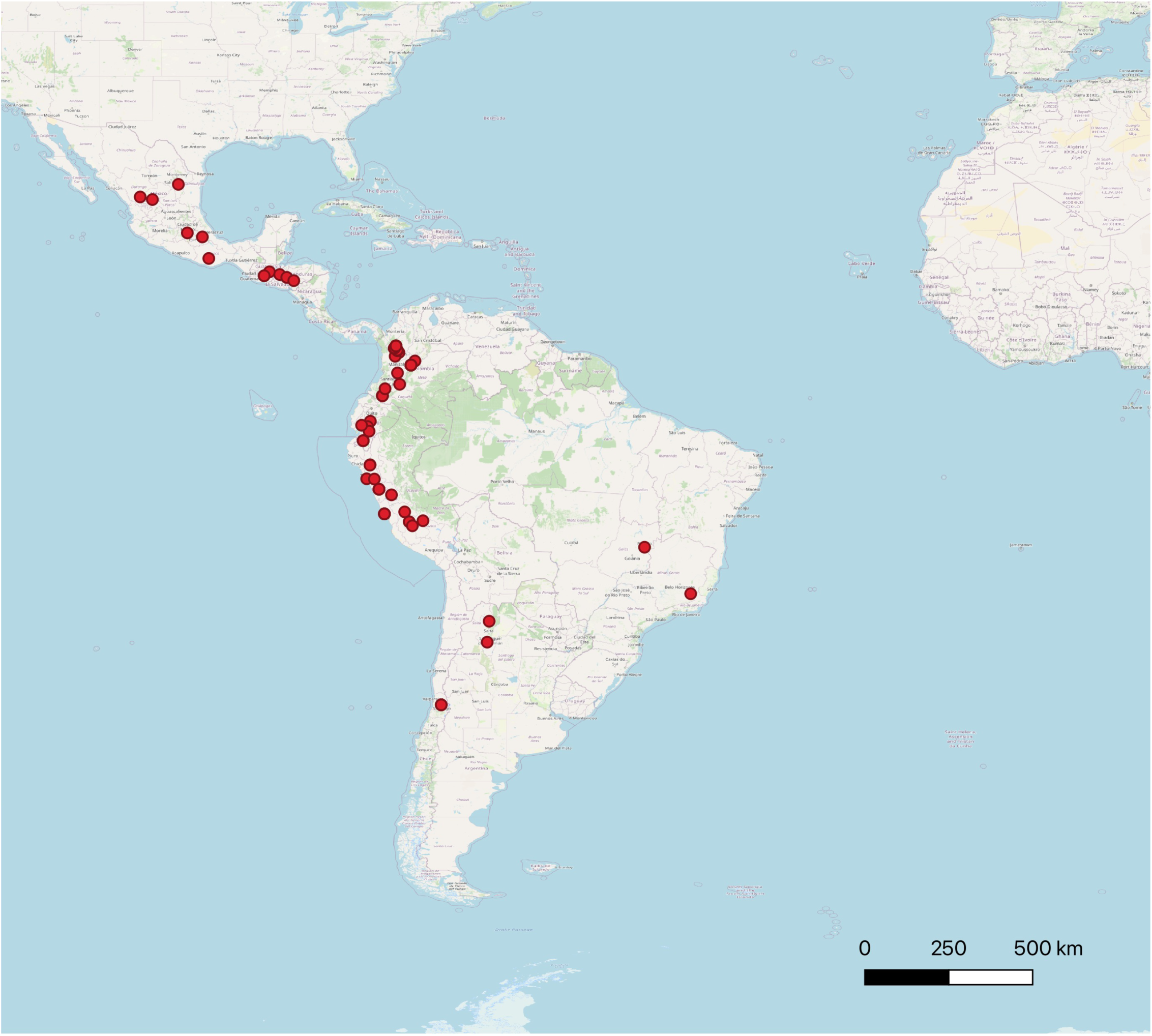
Distribution of the 127 common beans with location data that were used in this study. The coordinates of the capital city were used for those without coordinate data. Produced with QGIS.

### Genotyping

The genotypes were whole genome re-sequenced using Illumina short reads. The accessions were grown at the Norwich Research Park (Norwich, UK) in 2021 until the expansion of the first true leaf, after which they were snap-frozen (∼50-100 mg). The genomic DNA extraction for short read sequencing from each accession was completed using a Qiagen DNAeasy kit (Qiagen, Germany). The DNA concentration of the samples was quantified for quality control using the Tecan Plate Read Infinite F200 Pro for a fluorometry based assay. The sequencing of the samples was completed by Genomic services at Earlham Institute (Norwich, UK). LITE libraries, a cost-effective low volume variant of the standard Illumina TruSeq DNA protocol, were constructed for the 144 accessions and were sequenced using two NovaSeq 6000 S4 v 1.5 flow cells with 150bp paired-end reads, following the protocol in [47].

### Phenotyping

All 144 common bean accessions were evaluated at the Norwich Research Park (Norwich, UK) in temperature-controlled glasshouses. The experiments were conducted in two seasons; summer 2022 with long daylength (16:8) and winter 2023 with short daylength (12:12). The accessions were organised in a randomised block design with 3 or 2 replications, respectively. Management was conducted according to recommendations for common bean cultivation.

The diversity panel was characterised for the days to flowering (DTF), seed size (SS), weight of 100 seeds (E100_SW; estimated based on the weights of seeds harvested and projected to 100 seeds), determinacy (D; terminal flower bud presence) [25] and photoperiod sensitivity (PS; flowering in none, one or both seasons). DTF was split into the two seasons due to photoperiod sensitivity in certain accessions and PS was characterised in three ways for the GWAS.

The statistical analysis of variance (one-way ANOVA) of the phenotypic data was done in R, then the Pearsons’s correlation coefficient was calculated and visualised using the R package ‘corrplot’ [48].

### Pre-processing genotype data

The raw sequence reads were processed with TrimGalore (v. 0.5.0) [49] to remove adapters and poor-quality reads, and then quality checked using fastqc [50] and multiqc [51]. The trimmed reads were aligned to the Andean reference genome, *Phaseolus vulgaris* G19833, v2.1 [52] downloaded from Phytozome [53] with BWA-MEM (v 0.7.13) [54] and ‘-M -R’ to add read group information and allow compatibility with GATK. SAMtools (v 1.7) combined, compressed and sorted the aligned files [55]. Picardtools (https://broadinstitute.github.io/picard/) (v 2.1.1) marked duplicates and BamTools indexed the alignments [56]. The percentage of alignments were calculated at this stage. The genotype data was divided in 10 Mbp regions [57] (v 1.0.2) to run Genome Analysis ToolKit (GATK v 4.2) haplotype caller with default parameters [58]. This identified 20.2 million variant loci (∼17.1M SNPs and ∼3.4M indels).

### Population structure analysis

The resulting vcf file from GATK using the Andean reference (“Andean vcf”) was filtered further with BCFtools to retain calls with a minimum depth of 5 reads per variant call (FMT/DP >= 5), a locus call quality over 30, maximum missing calls per locus of 5%, to keep only biallelic SNP locus, and for a minor allele frequency over 2%. The resulting VCF had ∼9 million SNP loci. Then, the vcf was filtered for a maximum heterozygosity of 20% per locus using TASSEL 5 (v. 20230314) [59]. This was then filtered for linkage disequilibrium (LD) (based on LD decay) and thinned with a window size of 10 bps using BCFtools prune.

The population structure of the panel was analysed using ADMIXTURE (v 1.3.0) [60] on a subset of 88,786 SNP loci. ADMIXTURE was run for K=2 to K=10 and the ideal number of K was determined using the cross-validation error. Accessions were allocated a group when their membership coefficient (q) was greater than 0.7. Plotting was completed in R using the packages ‘ggplot2’ [61].

### Genome wide association study

The “Andean vcf” from GATK was filtered with BCFtools (v 1.12) [55] for biallelic loci, a minor allele frequency of 1% and thinned with a window size of 5 bp. To understand the genetic relationship between accessions we used a principal component analysis (PCA) generated with GAPIT v.3 [62] on a subset of 2,572,124 loci.

A genome-wide association study investigated marker-trait association for determinacy and photoperiod insensitivity phenotypes using GAPIT v.3 [62] with three principal components. We ran with the models Bayesian-information and Linkage-disequilibrium Iteratively Nested Keyway (BLINK) [63], Fixed and random model Circulating Probability Unification (FarmCPU) [64] and Mixed Linear Model (MLM) [65]. GAPIT was run on the whole panel (144 accessions) and on the Andean subpanel (as defined at K2 ADMIXTURE; 108 accessions). To run BLINK, GAPIT completed the analysis with the option “Random.model=TRUE” as to not calculate R^2^ for phenotypic variance explained (PVE) values after GWAS. The quantile-quantile (QQ) plots were used to understand the suitability of the models to the data. Plotting was completed in R using the package ‘ggplot2’ [61].

### Selecting significant loci, candidate gene mining and functional annotation

Significant marker-trait associations (MTAs) were investigated further when they had a - log_10_(p-value) over 7 and were confirmed by two models from GAPIT. QTLs were defined as ±100 kbp from the MTA [66, 67]. This is shorter than the calculated recombination rate in common bean of 3.72 cM/Mb [68].

Identified loci were compared to the Andean reference genome, *Phaseolus vulgaris* G19833 v2.1 in JBrowse [52, 69] whilst considering “high impact” mutations identified by SnpEff [70]. Once genes were identified, their putative function was explored using PhytoMine [53] (*Phaseolus vulgaris v.2*), BLAST [71] against the non-redundant (nr) protein database at NCBI, and finally against the TAIR database if no gene function could be identified in close relatives [72]. The loci were compared to previous studies and literature. PulseDB was used for comparison, particularly for QTLs and markers related to developmental and flowering phenotypes [73]. QTLs and markers were mapped to the reference genome to estimate the conversion from cM to Mb in JBrowse.

## Results

### Population structure

The diversity panel split into the two gene pools, the Andean and Mesoamerican (figure 2A, figure 3A). At K6 (figure 2B), the Mesoamerican group split into two subpopulations (M1 and M2), while the Andean subgroup split into 4 subpopulations. Two of these subpopulations included only accessions from Colombia and were named C1 and C2. A subpopulation containing accessions from Colombia, Ecuador/Peru was named C-EP. The remaining subpopulation was named A1. In the PCA (fig. 3A), PC 1 explained 38.8% of the variation in our diversity splitting the two gene pools, while PC2 accounted for 5.06% of the variation, splitting the Mesoamerican subgroups (M1 and M2) and separating C-EP from the other Andean subgroups. A total of 11 accessions were classified as admixed between the Andean and Mesoamerican gene pools (Admx_AM), as they had an ancestry composition lower than 70% from either of the origins (q<0.7). The Admx_AM accessions were all indeterminate and produced a variety of seed sizes. Seven were landraces and two were wild. There was also a mix of photoperiod sensitive and insensitive accessions.

**Figure 2:**
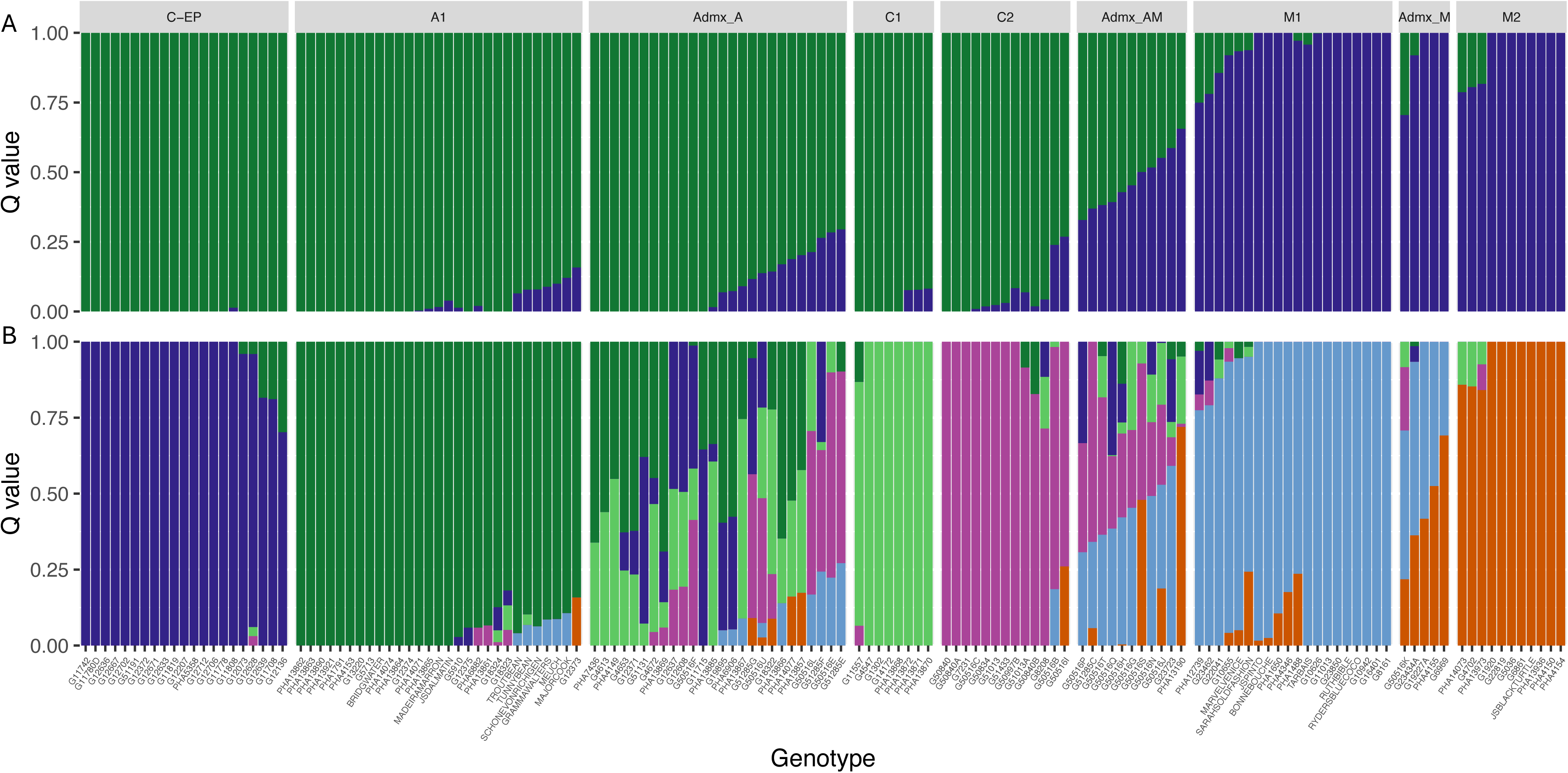
Analysis of the population structure of 144 accessions belonging to our diversity panel focusing on Colombia at K=2, Andean or Mesoamerican groups (A) and K=6 (B). (C-EP) accessions mainly from Peru, then Ecuador and Colombia; (A1) Andean accessions from a variety of South American countries; (C1) mostly determinate Colombian landraces; (C2) indeterminate Colombian landraces; (M1) mainly medium seeded** from Central America and Colombia; (M2) mainly small seeded** from Central America and Colombia. (Admx_AM) Andean X Mesoamerican hybrids; (Admx_A) and (Admx_M) admixed accessions between subpopulations (ancestry composition q <0.7 at K=6). ** p < 0.01 using a two tailed student t-test with unequal variance.

**Figure 3:**
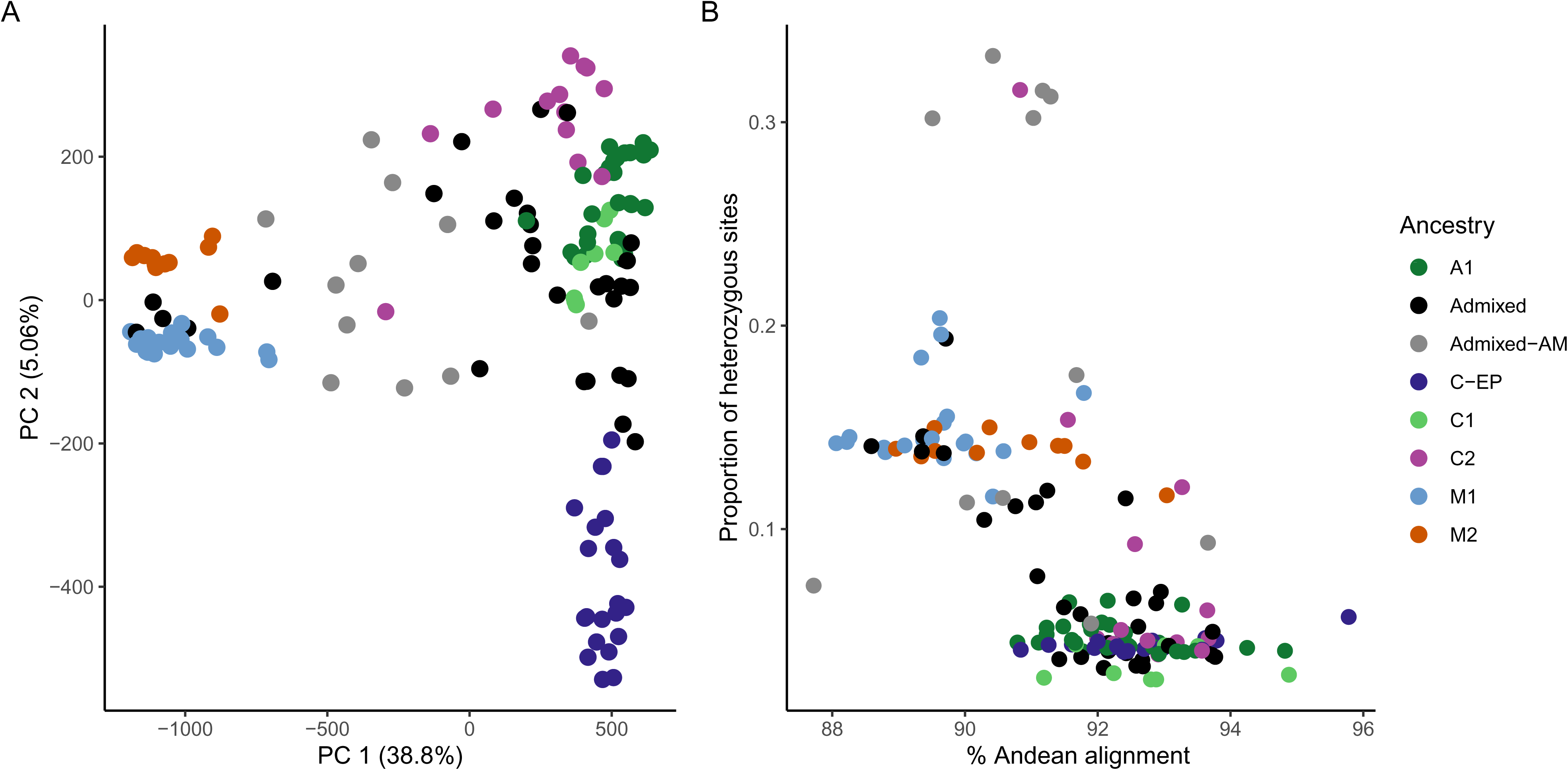
(A) Principle component analysis (PCA) plot of PC1 against PC2. (B) Proportion of heterozygous sites against the percentage of read-pair alignment to the Andean reference genome G19833 [52]. The colours illustrate the population structure of our diversity panel.

The Colombian subgroups (C1 and C2; figure 2B) contained medium and large seeded landraces. However, the subpopulations distinguished by determinacy; C1 contained mainly insensitive determinate accessions while C2 contained sensitive indeterminate accessions. The A1 group contained large and medium seeded landraces that were mainly photoperiod insensitive. The C-EP population contained accessions from Ecuador, Peru and Colombia. This group contained large-seeded indeterminate landraces and also included accessions from races previously identified to be from the Andean gene pool. The Mesoamerican subgroups (M1 and M2; figure 2B) were also distinguished by phenotypic data. They both contained indeterminate and determinate accessions; however, M1 were mainly medium seeded while M2 were mainly small seeded. This is summarised in Table 1 and supplementary table 1.

**TABLE 1:** Phenotypic characteristics associated with each subpopulation.

| **Subpopulation** | **Gene pool** | **Determinancy** | **Photo. Sen.** | **Seed size** | **Origin** |
| --- | --- | --- | --- | --- | --- |
| **C1** | Andean | Mainly determinate | Insensitive | Mainly large | Colombia and Ecuador |
| **C2** | Andean | Indeterminate | Mainly sensitive | Mainly large | Colombia |
| **A1** | Andean | Both | Mainly insensitive | Mainly large | South America, Heirlooms, Colombia |
| **C-EP** | Andean | Indeterminate | Sensitive | Large | Colombia, Ecuador, Peru |
| **Admix_A** | Andean | Mainly indeterminate | Both | Mainly large | Colombia and South America |
| **M1** | Mesoamerican | Mainly indeterminate | Both | Mainly medium** | Central America, Colombia, Heirlooms, Peru |
| **M2** | Mesoamerican | Mainly indeterminate | Mainly insensitive | Mainly small** | Central America, Colombia |
| **Admix_M** | Mesoamerican | Mainly indeterminate | Insensitive | Small and medium | Colombia, Brazil, Heirlooms, Central America |
| **Admix_AM** | AxM hybrids | Indeterminate | Mainly sensitive | Mainly medium | Colombia and Ecuador |

Colombian accessions can be found within all the subgroups and admixed groups at K=6 (figure 2B). While the admixture accessions are mainly from Colombia, while one sample is a wild “Ecuador” accession.

The Andean accessions had a lower proportion of heterozygous sites (<0.1) than the Mesoamerican accessions, which were more heterozygous (fig. 3B). The six highly heterozygous accessions (>25% of the loci) were found within the Andean X Mesoamerican hybrid (Admixed-AM) subpopulation (Figure 3B) and were from Colombia. Finally, the outlier accession with the lowest alignment to the Andean reference genome and low proportion of heterozygous sites was a wild accession from Ecuador.

### Phenotypic variation and correlations

The correlation coefficient was estimated for each pair of traits (fig. 4), averaged over two seasons or studied in both years. There was a positive correlation between DTF from winter and summer (r=0.57). Both DTF were negatively correlated with PS (r=−0.72 (DTF_S22), r=−0.77 (DTF_W23)) and D (r=−0.35 (DTF_S22), r=−0.43 (DTF_W23)). Population structure at either two or six ancestries (K2, K6) was positively correlated with D (r= 0.32 (K6), r= 0.37 (K2)) but negatively correlated with SS (r= −0.44 (K6), r= −0.4 (K2)) and E100_SW (r= −0.37 (K6), r= −0.47 (K2)). SS was not correlated with DTF_S22, DTF_W23, D or PS (r= −0.13, r= - 0.07, r= −0.12, r= 0.09). However, E100_SW was positively correlated with PS (r= 0.18) and SS (r=0.87) but negatively correlated with DTF_S22 (r= −0.22). Then D and PS were positively correlated (r=0.45).

**Figure 4:**
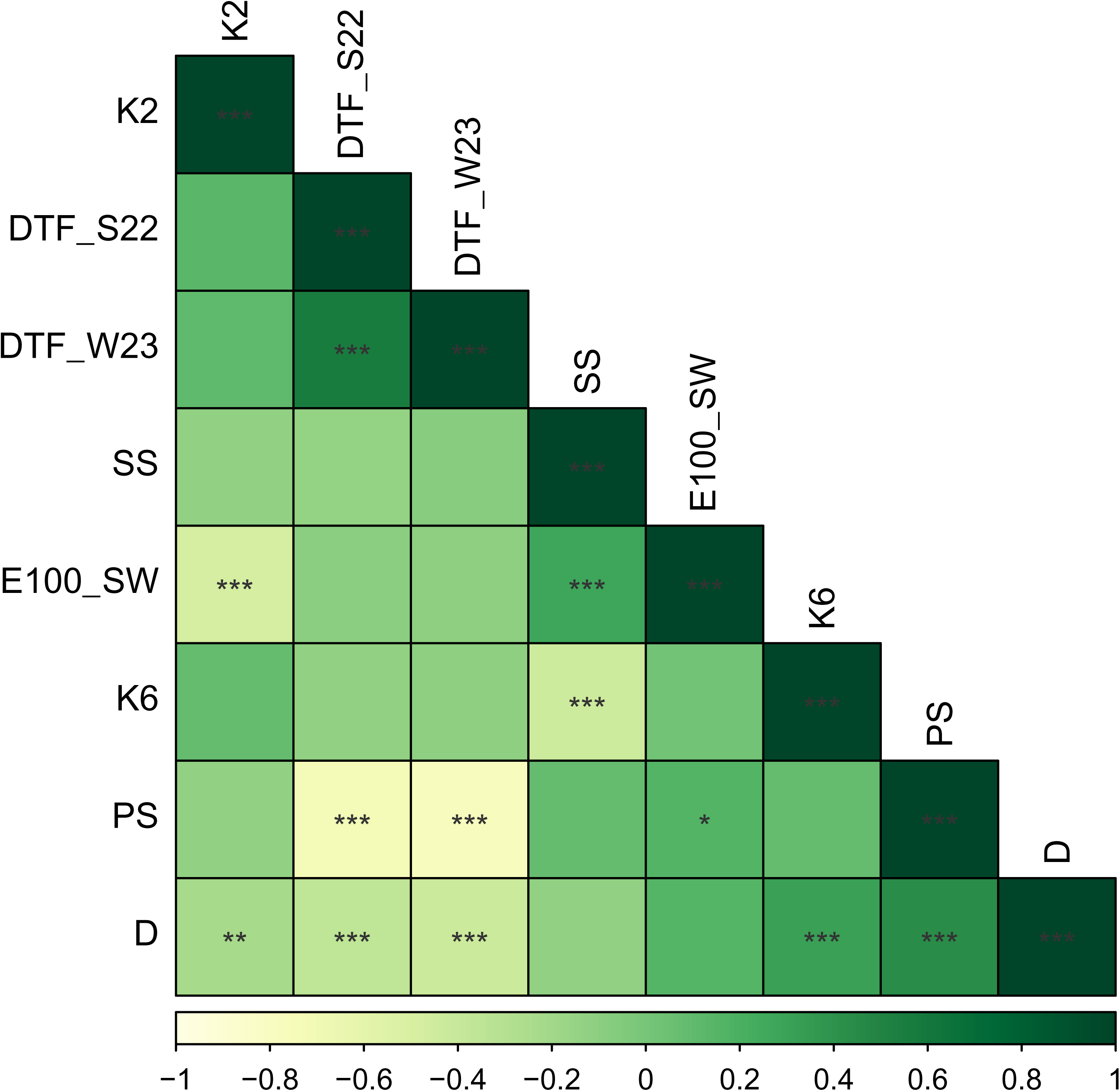
Pearson correlation coefficients among 5 agronomic traits and population structure measured in 144 common bean genotypes grown at the Norwich Research Park, Norwich, UK in 2022 and 2023. (K6) K6 subgroups from ADMIXTURE; (K2) K2 subgroups from ADMIXTURE; (D) Determinacy; (PS) photoperiod sensitivity; (SS) seed size; (E100_SW) Estimated weight of 100 seeds; (DTF_W23) days to flowering from winter 2023; (DTF_S22) days to flowering from summer 2022. *Indicates p < 0.05; **indicates p < 0.01; ***indicates p < 0.001.

Figure 5A, 5B and 5C showed the distributions of the phenotyping for traits E100_SW, S22_DTF and W23_DTF, respectively. The seed weights (fig. 5A) were normally distributed, while the DTF in summer and winter (figs. 5B and 5C) were binomial distributions; the peaks were around 42- and 54-days post-sowing in summer, and around 70- and 90-days in winter. When analysing the phenotypes by subpopulation, we can see that C-EP (figure 2B) did not flower during winter in the UK, W23_DTF, as were mainly photoperiod sensitive. This is further supported by the correlation plot (Figure 4). Furthermore, determinacy, photoperiod insensitivity and days to flowering are correlated. The determinate accessions flower earlier than the indeterminate, supporting the binomial distribution.

**Figure 5:**
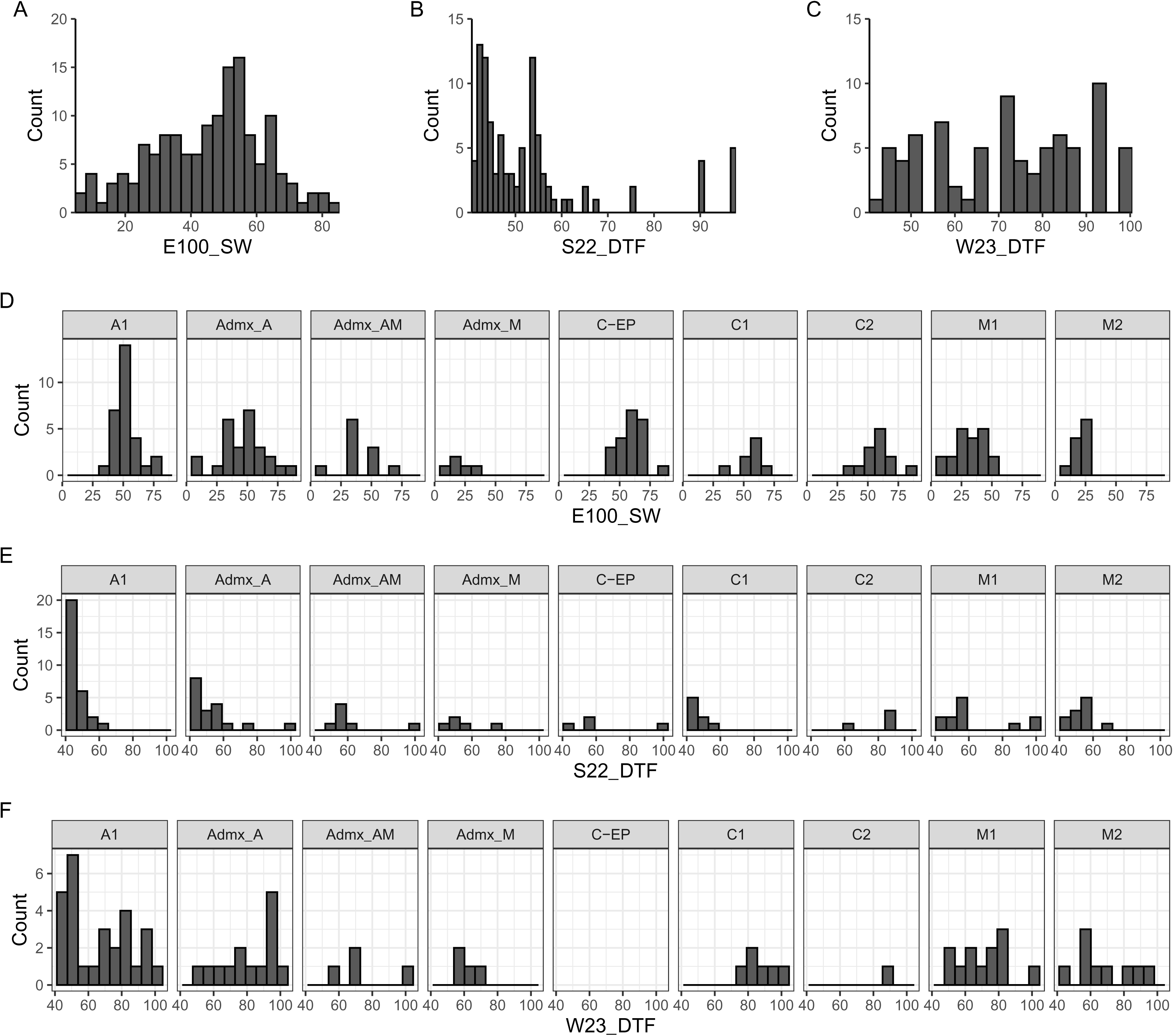
Frequency distribution of seed weight and days to flower traits evaluated in two seasons in a common bean diversity panel. (A) E100_SW, Estimated weight of 100 seeds; (B) phenological days to flowering in the summer 2022 (S22_DTF) (and (C) in the winter 2023 (W23_DTF) at the Norwich Research Park, excluding those which didn’t flower. The distributions were split into the subpopulations from K6 ADMIXTURE. (D) E100_SW***; (E) S22_DTF***; (F) W23_DTF*. Completed a one-way ANOVA for E100_SW, S22_DTF and W23_DTF. *Indicates p < 0.05; **indicates p < 0.01; ***indicates p < 0.001.

### GWAS for Determinacy

The GWAS was performed using the models BLINK, FarmCPU and MLM with GAPIT (figures 6A and 6B). The QQ plots (figures 6C and 6D) provided evidence that the selected models were well fitted to identify significant marker trait associations (MTAs) for the dataset. We identified 13 MTAs with a significant p-value, corresponding to 13 QTLs. We focused on seven significant MTAs that were identified for the whole panel based on the criteria laid out in the methods (vertical lines in figure 6). The seven QTLs were found on chromosomes Pv01, Pv07, Pv08, Pv09 and Pv10 (Table 2). Five of the seven QTLS were also identified for the Andean subset.

**Figure 6:**
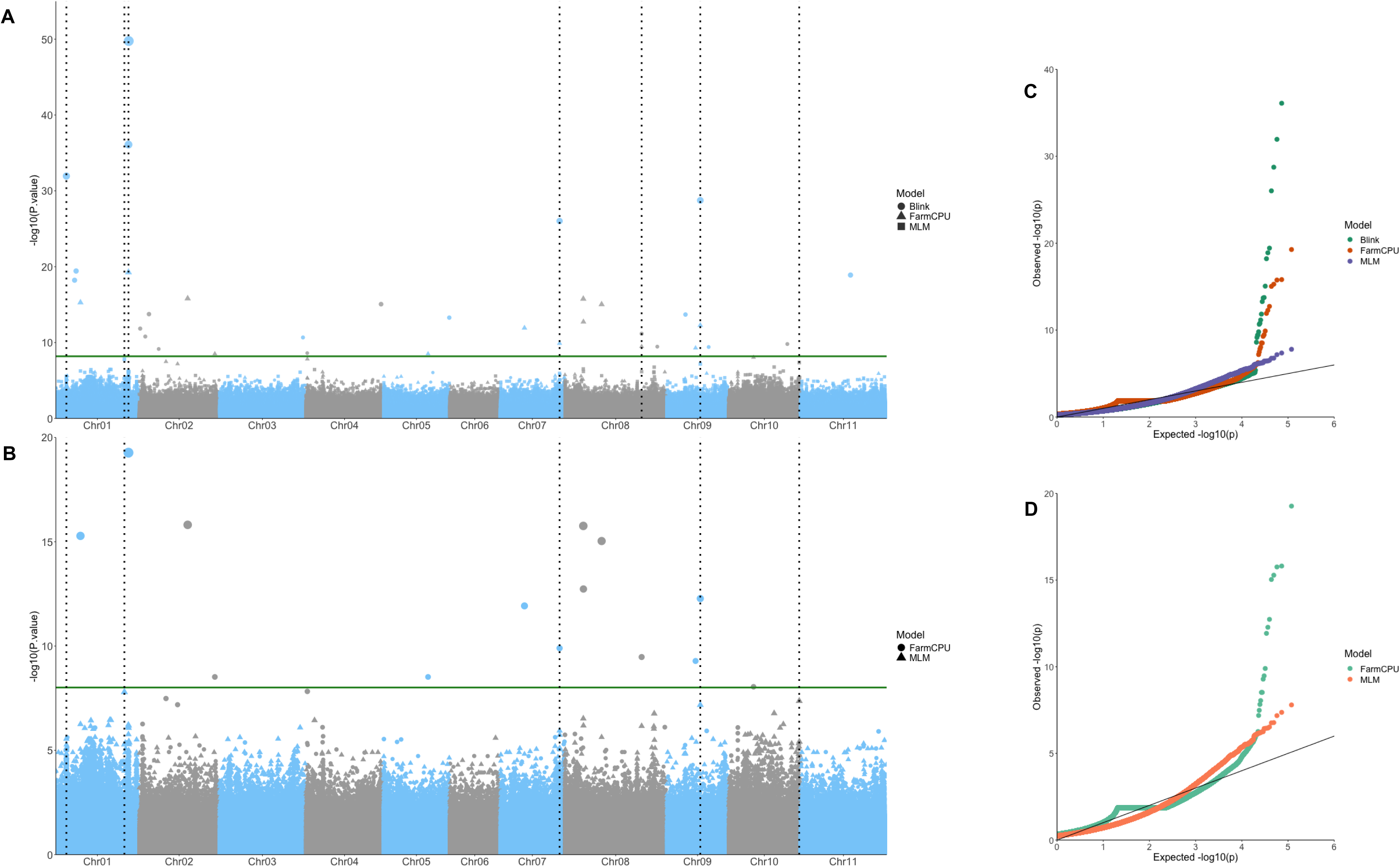
Manhattan plots highlighting markers significantly associated with determinacy on (A) the whole panel and (B) the Andean subpanel. The analyses were completed with GAPIT and the models FarmCPU, BLINK or MLM [62–65]. The X-axis represents the genomic position of markers and the Y-axis is the -log 10 of the P-values for association with the phenotype. The vertical lines correspond to QTLs found by at least two models. Point size correlates to -log10(P-value). Quantile-quantile (QQ) plots are provided for (C) the whole panel and (D) the Andean panel.

**Table 2:** QTLs for determinacy and photoperiod sensitivity.

| Name | Chromosome | Start | End | Trait | Panel |
| --- | --- | --- | --- | --- | --- |
| D1.1 | Chr01 | 6,512,000 | 6,521,000 | Determinacy | Andean + Whole |
| D1.2 | Chr01 | 11,363,000 | 11,372,000 | Determinacy | Andean |
| D1.3 | Chr01 | 42,404,000 | 42,413,000 | Determinacy | Andean + Whole |
| D1.4 | Chr01 | 44,856,000 | 44,847,000 | Determinacy | Whole |
| D1.5 | Chr01 | 44,932,000 | 44,941,000 | Determinacy | Andean + Whole |
| D1.6 | Chr01 | 45,098,000 | 45,107,000 | Determinacy | Whole |
| D2.1 | Chr02 | 24,821,000 | 24,830,000 | Determinacy | Andean |
| D3.1 | Chr03 | 25,608,000 | 25,617,000 | Determinacy | Andean |
| PS4.1 | Chr04 | 38,316,000 | 38,325,000 | photo sens | Whole |
| PS5.1 | Chr05 | 16,423,000 | 16,432,000 | photo sens | Whole |
| PS5.2 | Chr05 | 18,321,000 | 18,330,000 | photo sens | Andean |
| PS7.1 | Chr07 | 16,829,000 | 16,838,000 | photo sens | Andean + Whole |
| PS7.2 | Chr07 | 26,485,000 | 26,494,000 | photo sens | Andean + Whole |
| D7.1 | Chr07 | 36,860,000 | 36,869,000 | Determinacy | Andean + Whole |
| PS8.1 | Chr08 | 4,234,000 | 4,243,000 | photo sens | Whole |
| D8.1 | Chr08 | 7,440,000 | 7,449,000 | Determinacy | Andean |
| PS8.2 | Chr08 | 8,320,000 | 8,329,000 | photo sens | Andean |
| D8.2 | Chr08 | 47,582,000 | 47,591,000 | Determinacy | Whole |
| D9.1 | Chr09 | 20,814,000 | 20,823,000 | Determinacy | Andean + Whole |
| PS9.1 | Chr09 | 21,640,000 | 21,649,000 | photo sens | Whole |
| PS9.2 | Chr09 | 34,445,000 | 34,454,000 | photo sens | Andean |
| D10.1 | Chr10 | 43,762,000 | 43,771,000 | Determinacy | Andean + Whole |
| PS11.1 | Chr11 | 204,000 | 213,000 | photo sens | Andean |

Putative candidate genes were identified for determinacy based on the significant MTAs and corresponding QTL windows. The identified genes and QTLs are listed in supplementary tables 2 and 3.

### GWAS for Photoperiod sensitivity

The GWAS was performed using the BLINK and FarmCPU models with GAPIT (figure 7A and 7B). The QQ plots (figure 7C and 7D) provide evidence that the selected models are fitted to identify significant marker trait associations (MTAs) for the dataset. We identified 10 QTLs. We focused on six QTLs for the whole panel based on criteria laid out in the methods. The MTAs were found on chromosomes Pv04, Pv05, Pv07, Pv08 and Pv09 (vertical lines in fig 7). Six QTLs were identified for the Andean subset panel in Chromosomes Pv05, Pv07, Pv08, Pv09 and Pv11. The QTL in Pv04 and Pv09 were found in the full dataset only. The QTL in Pv9 and Pv11 were found in the Andean subset only. Candidate genes were identified for the significant MTAs and their corresponding QTLs. The identified genes and QTLs are listed in supplementary table 2 and 3.

**Figure 7:**
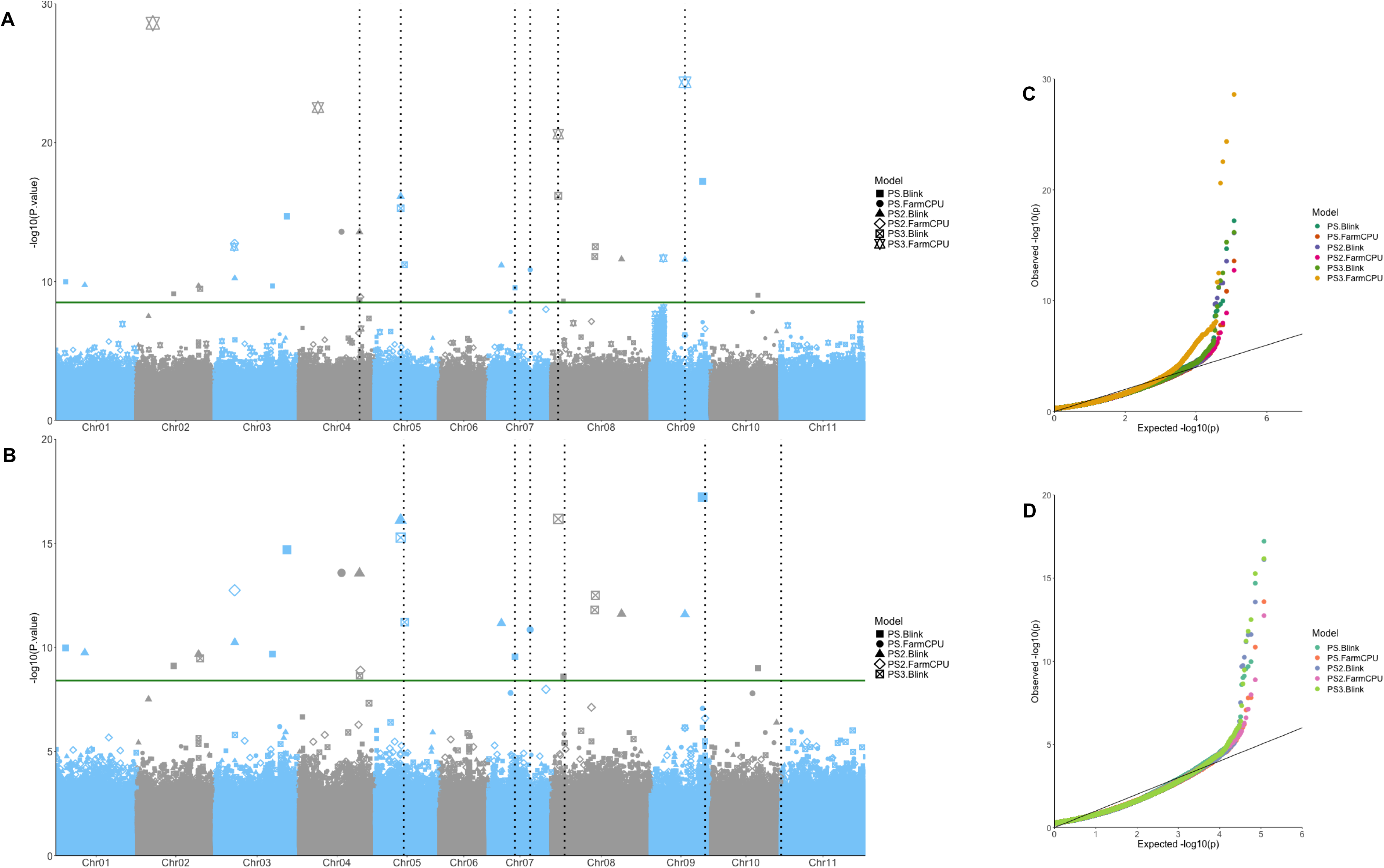
Manhattan plots highlighting markers significantly associated with photoperiod insensitivity on (A) the whole panel and (B) the Andean subpanel. The analyses were completed with GAPIT and the models FarmCPU, BLINK or MLM [62–65]. The X-axis represents the genomic position of markers and the Y-axis is the -log 10 of the P-values for association with the phenotype. The vertical lines correspond to QTLs found by at least two models. Point size correlates to -log10(P-value). Quantile-quantile (QQ) plots are provided for (C) the whole panel and (D) the Andean panel.

## Discussion

We delimited subpopulations in a panel of 144 accessions, initially divided by domestication event into the two Andean and the Mesoamerican gene pools (Figure 2, Figure 3) [42, 74]. The Mesoamerican gene pool is generally more diverse [52, 75] with less influence from domestication bottlenecks. Furthermore, the Mesoamerican gene pool within our diversity panel is also more heterozygous, suggesting that the Andean gene pool has undergone fewer outcrossing events [76]. However, care needs to be taken when utilising market sampling information. This is highlighted by the two ‘Peruvian’ accessions collected from markets that fall with the Mesoamerican subpopulation (supplementary table 1).

Admixture was commonly observed in the panel, including 26 admixed Andean accessions, 5 admixed Mesoamerican accessions, and 11 Mesoamerican x Andean accessions. This supports our initial hypothesis that Colombia and neighbouring countries hold large common bean variation, including hybrids between both gene pools [77–79]. The wider crosses between gene pools compared to within gene pools resulted in a larger observed heterozygosity in the hybrid accessions, supporting the outcrossing events and movement between gene pools. One implication of this study is that admixed Colombian hybrid landraces bridge Andean and Mesoamerican gene pools, and novel allelic and epistatic interactions likely filtered out deleterious effects [80] due to stronger purifying selection with increased recombination. After all, recombination increases local effective population size (Ne) and limits Hill-Robertson interference [81]. This suggests the Colombian hybrids have promising potential for breeding. However, the diversity panel may be also biased and underestimating their prevalence in other regions due to the large number of Colombian accessions in our diversity panel.

We observed some traits associated with demography, including determinacy and photoperiod sensitivity: C1 and C2 shared origin but could be separated by ancestry admixture analysis, and were characterised by different determinacy, as C1 contained mainly determinate accessions, and C2 mainly indeterminate accessions. Furthermore, the population structure suggests that Colombian farmers have not selected varieties based on the seed characteristics studied (e.g. seed size) [82].

Indeterminate and photoperiod sensitive landraces were common, despite the combined selection for photoperiod insensitivity and determinacy resulting in common bean varieties with shorter flowering periods (DTF) and easier management. Prior research supports the correlation between DTF and phenotypes such as seed weight, determinacy and growth habit [66, 76, 83–85]. These phenotypes are related to apical meristems and floral development [86].

We observed the distribution of DTF values, in either summer or winter, were bimodal, i.e. had two peaks (figure 5B and 5C). This likely occurred due to the determinate types flowering first, then followed by the indeterminate beans [87]. The distribution also correlates to growth habits as bush types typically flower earlier than climbing types [88]. Figure 2A supports that photoperiod sensitivity arose during domestication in both gene pools [24].

The Andean accessions within our diversity panel were large and medium seeded while the Mesoamerican accessions were small and medium sized, which supports previous research [89]. Among the Mesoamerican accessions, the Mesoamerican race is characterised by small-seeds, while the race Durango-Jalisco (DJ) is characterised by medium-seeds [89–92]. We could not separate our diversity panel into subpopulations matching these races due to a lack of Mesoamerican diversity in the panel, a limited genetic component for the seed size trait, or introgressions occurring in the Mesoamerican Colombian accessions.

Interestingly, Ecuador accessions are often separated from Andean subgroups, suggesting that they are members of the PhI group or a possible sister species *Phaseolus debouckii* [93, 94]. Further to this, the wild Ecuador accession is separated from both gene pools (Figure 2 and 3), suggesting a separate ancestry originating from Ecuador or Peru [19, 95]. Finally, the C-EP group (figure 2B) are mainly photoperiod sensitive (figure 5F), possibly due to a different domestication history or due to their Equatorial provenance not necessitating evolution under fluctuating photoperiods.

By leveraging this diversity panel and its trait segregation across the demographic stratification, we prioritised 13 QTLs for determinacy and 10 QTLs for photoperiod sensitivity. Four of the QTLs for photoperiod sensitivity, and four for determinacy, were also identified only for the Andean subset, but not the whole panel. The Andean gene pool has adapted to lower latitudes than the Mesoamerican pool, resulting in differential selection for photoperiod sensitivity between the two gene pools.

## QTLs and candidate genes associated with determinacy

### Three QTLs in Chromosome 1

We identified a determinacy QTL in chr 1 -Pv01-(D1.4-D1.6; table 2), identified in other studies [66, 83, 96–99] as a hotspot of allelic variation, named the *Fin* locus. The *Fin* locus has been mapped to ∼44.5Mb [98, 100]. This co-segregates with an upstream gene, *TFL1y* (*Phvul*.001G189200), a candidate gene for flowering, vegetative growth, rate of plant production and determinacy [23, 26, 28, 44, 67, 80, 101]. Consequently, the *Fin* locus has pleiotropic effects due to associations with many development traits such as determinacy, shoot biomass, days to flowering, days to maturity, plant architecture, embryo abortion, number of pods per plant, and number of seeds per plant (seed yield and weight) [28, 101]. However, segregation for this QTL hotspot in Pv01 may prove difficult in breeding programmes due to these pleiotropic effects [83].

Further candidate genes have been identified in this QTL, such as *Phvul.001G192200*. This gene is an ortholog of *LIGHT-REGULATED WD1* (*LWD1*), a gene involved in the circadian rhythm pathway [66, 101, 102], or *Phvul.001G192300,* which is an ortholog of *SPINDLY* (*SPY*). *SPY* interacts with genes in the reproductive pathway [66, 97, 103] and has been associated with days to maturity [104].

Another QTL we identified on Pv01 (D1.3; table 2) contains the gene *Phvul.001G168700*. This gene is related to the PIF1 (Phytochrome interacting factor1) transcription factor isoform X1 in the legume *Vigna radiata* [105]. This bHLH transcription factor is involved in many light dependent pathways in plant development and interacts with circadian clock genes [106].

### QTL D7.1 in Chromosome 7

The QTL at Pv07 (D7.1) was identified in the whole and Andean panel. The QTL contains the gene *Phvul.007G244700.* This is related to a transcriptional corepressor, Leunig-homolog (LUH) in *Vigna radiata* [105]. In *Arabidopsis* Luenig-homologs have functional redundancy with Leunigs (LUGs), and are involved in embryo and floral development [107]. This QTL has been associated with seed size, seed weight and growth habit [44, 84, 96, 97], suggesting it may have pleiotropic effects.

### QTL D8.2 in Chromosome 8

The QTL identified on Pv08 (D8.2; table 2) for determinacy has previously been identified for plant architecture [97]. However, no gene with a clear function was identified. We have, however, identified a possible candidate gene for further investigation; *Phvul.008G170000.* This encodes a putative FAF (fantastic four) domain-containing protein. In *Arabidopsis,* FAF proteins regulate shoot meristem size and architecture [108].

### QTL D9.1 in Chromosome 9

The QTL D9.1 in chr 9 was identified in the whole and Andean panel. Nearby QTLs have been identified for yield and determinacy [67, 109]. The gene *Phvul.009G138100* is found within this QTL and contains the significant MTA found by GAPIT [62]. This gene has an insertion that possibly affects function [70]. This gene is uncharacterised in common bean but has homology to the Root Meristem growth factor 9 from *Glycine soja* [53, 105]. This growth factor is expressed in the roots and flowers, regulating and maintaining apical meristems, and therefore both root and floral development, seed size and leaf architecture [110, 111]. Although has previously identified as a candidate gene associated with Mesoamerican domestication [52], we found the QTL in the Andean panel, suggesting that it has also played a role in the Andean domestication event.

### QTL D10.1 in Chromosome 10

The QTL on Pv10 (D10.1) is located near to QTLs for plant height and number of nodules, once again suggesting pleiotropic effects [101]. Three of the genes within this region encode bHLHLZip proteins: *Phvul.010G158500, Phvul.010G158300* and *Phvul.010G158200.* These bHLH transcription factors may be involved in the regulation of flowering genes [112]. The gene *Phvul.010G158500* displays non-synonymous modifications in our panel, including insertions, deletions and other variants linked to frameshift mutations and gained stop codons [70]. Homology to *Vigna angularis* suggests this gene may be related to the transcription factor bHLH25, and possibly linked to a circadian rhythm-associated protein [53].

## Candidate genes for photoperiod sensitivity

### QTL PS4.1 in Chromosome 4

One QTL for photoperiod sensitivity was found on Pv04 (PS4.1; table 2) from the analysis on the whole panel. Within this QTL, four genes were identified, three of which (*Phvul.004G110200*, *Phvul.004G110301*, *Phvul.004G110000*) have non-synonymous mutations such as a stop lost, stop gained or a frameshift mutation in our panel [70]. However, the genes are uncharacterised.

### Two QTLs in Chromosome 5

Two QTLs were identified in Pv05: PS5.2 for the Andean panel and PS5.1 for the whole panel. PS5.2 overlaps with a previously identified QTL for seed weight, days to flowering and pod weight [104, 113]. However, this previous analysis with a limited number of markers did not identify a candidate gene. Based on sequence homology with *Vigna radiata,* we identified the gene *Phvul.005G077000*, which encodes a Proton gradient regulation 5 (PGR5) protein [105]. PGR5 is involved in plant growth under different light conditions due to interactions with Photosystem I, and consequently putatively associated with differentiating photoperiod sensitivity in our panel [114]. The QTL PS5.1 contained two genes, one of which, *Phvul.005G076300*,may encode a bidirectional sugar transporter, named SWEET protein. Evidence suggests SWEET proteins have essential roles in plant development, including in reproductive organs and bud growth [115].

### Two QTLs in Chromosome 7

Two QTLs were also identified on Pv07. PS7.1 and PS7.2, both in the Andean and the whole panel. The QTL PS7.2 contains the genes *Phvul.007G157400* and *Phvul.007G156200*. Homology with *Arabidopsis* suggests that *Phvul.007G157400* encodes a BANQUE3 BHLH161 protein. BANQUE3 is negatively regulated by *APETALA3* and *PISTILLATA* in petals and is involved in light-regulated responses and flowering time [72, 116]. *Phvul.007G156200* may encode the BHLH transcription factor PIF4 (Phytochrome Interacting Factor 4) based on homology with *Vigna radiata* and *Glycine soja* [53, 105]. PIF4 is a downstream signalling component integrating environmental cues such as light [105].

The QTL PS7.1 overlaps with a previously identified QTL for plant production traits [28]. The QTL includes the gene *Phvul.007G117400* which encodes a putative JUMONJI domain containing protein [53]. JUMONJI proteins are involved in multiple plant developmental processes such as flowering and leaf senescence [117, 118]. *Phvul.007G117400*’s homology with a JUMONJI16 orthologue in *Vigna radiata* also supports this role [105].

### Two QTLs in Chromosome 8

One of the QTLs found in Pv08 is PS8.1 from the whole panel. This QTL has been associated with determinacy [67], seed weight [84], days to flowering [119] and pod number [109]. Due to the marker technology used, the QTL for seed weight was large so had low resolution [84]. Our results (figure 4) suggest a correlation between days to flowering, determinacy and photoperiod sensitivity under the same QTL. The significant MTA for this QTL was within the gene *Phvul.008G048300*. However, the function of this gene is currently unclear.

The other QTL found on Pv08 is PS8.2, which has previously been identified for seed weight [120]. Genes within this QTL include *Phvul.008G085000, Phvul.008G084500, Phvul.008G084900* and *Phvul.008G084100*. *Phvul.008G085000* is homologous to *gibberellin 2-oxidase 8* in *Arabidopsis* [72]. Gibberellin oxidases may respond to light intensity, and can therefore be related to photoperiod sensitivity [121]. *Phvul.008G084100* is homologous to *CLAVATA3* in *Arabidopsis, a* gene that regulates shoot and floral meristem development [122, 123]. *Phvul.008G084900* is homologous to genes encoding ovate family proteins (OFPs). OFPs appear to be sensitive to light stimuli [124]. *Phvul.008G084500* has homology with *RAVEN/INDETERMINATE DOMAIN5* in *Arabidopsis,* which is linked to GA signalling pathways as well as other plant developmental pathways [125, 126]. *Phvul.008G085000* and *Phvul.008G084900* also both contain insertions or deletions with high impact non-synonymous mutations which, therefore, possibly affect function [70].

### Two QTLs in Chromosome 9

A QTL was identified on Pv09 in the Andean panel (PS9.1). This was near a QTL associated with grain yield [84], post-harvest index [99], shoot biomass [98], seed size [97], days to flowering, and yield [120]. Genes within the QTL included *Phvul.009G229100, Phvul.009G229200, Phvul.009G229700* and *Phvul.009G229900. Phvul.009G229100* is homologous to PIN3 transcription factor genes, involved in regulating root and shoot growth [53, 127]. Homology with *Arabidopsis* suggests *Phvul.009G229200* and *Phvul.009G229700* are involved in root growth [72], and that *Phvul.009G229900* encodes a *HAB1 (Hypersensitive To Aba1) homology to ABI (Abscisic Acid-Insensitive)1* gene involved in ABA signal transduction, which is regulated by circadian rhythm [128, 129]. The other QTL in PV09 (PS9.2) was found in the whole panel and included the gene *Phvul.009G145100*, which was also related to an ABA response gene in *Arabidopsis*. A nearby QTL to PS9.2 was previously identified for days to flowering [96].

### QTL PS11.1 in Chromosome 11

The QTL at PV11 (PS11.1) was near a QTL for seed weight [97]. This may be due to pleiotropic effects or low resolution of the previous analysis with a limited number of markers. Within this QTL is the gene *Phvul.011G004000* which encodes a putative PHD finger protein. PHDs have been found to be involved in the regulation of flowering time [112, 130]. Other genes within the QTL are related to root or shoot growth. For example, homology of *Phvul.011G003200* and *Phvul.011G003400* implicates them in processes involved in root meristem development [72]. *Phvul.011G003700* is an uncharacterised gene in common bean but homology with *Arabidopsis* suggests it may be associated with phytochrome interacting factor 7 (PIF7) to regulate hypocotyl elongation [72, 131]. However, there are many genes within this QTL and further research is needed to clearly distinguish a candidate gene.

## Conclusion

Gene function analysis has relied on both forward and reverse genetic approaches in the model angiosperm Arabidopsis or the well-researched legume oil seed cash crop soybean. Current functional inferences, therefore, rely heavily on gene function information from either Arabidopsis or soybean and occasionally the more closely related Vigna bean species. As technologies improve, we expect common beans to be more easily analysed using reverse genetics to confirm the identity and function of genes. By linking candidate genes to phenotypes, we hope more targeted precision breeding approaches can be adopted to improve common bean traits under climate change. Nevertheless, this current study and previous ones highlight that for some genes and genomic regions, this will be difficult due to the high proportion of pleiotropic effects in common beans.

Furthermore, our research shows that common bean accessions from Colombia contain introgressive hybridisation and admixture diversity from the Andean and Mesoamerican gene pools. There was a systematic association between the population structure and agronomic traits such as determinacy and photoperiod sensitivity. Also, the identified genomic regions are connected to putative candidate genes involved in developmental and reproductive pathways. Consequently, GWAS are important in identifying MTAs and candidate genes, especially when accounting for population structure.

## Data availability

We thank CIAT’s Genebank and IPK’s Genebank for their generous provision of germplasm. Germplasm held in the CIAT and IPK collections is available on request. Raw reads are deposited in the SRA under accession PRJEB81566. The scripts used in this study are publicly available in Github (https://github.com/DeVegaGroup/KDJ-CBeans/).

## Acknowledgements

The authors declare that they have no conflict of interest. All the authors contributed and approved this manuscript. KDJ is supported by the Biotechnology and Biological Sciences Research Council (UKRI-BBSRC) through the Norwich Research Park Doctoral Training programme (#2578607). This research was partially funded by the British Council throughout the 2019 Newton Fund Institutional Links binational Bioeconomy grant ID 527023146 to AJC, JDV and APC. This study was also partially funded by the Biotechnology and Biological Sciences Research Council (BBSRC), part of UK Research and Innovation (UKRI), via Earlham Institute’s Strategic Programme Grant “Decoding Biodiversity” (BBX011089/1), and its constituent work package BBS/E/ER/230002B (Decode WP2 Genome Enabled Analysis of Diversity to Identify Gene Function, Biosynthetic Pathways, and Variation in Agri/Aquacultural Traits). Funding was also received from the BBSRC Core Strategic Programme Grant (Genomes to Food Security) BB/CSP1720/1 and its constituent work package BBS/E/T/000PR9818 (WP1 Signatures of Domestication and Adaptation), as well as BB/X011089/1. The authors would like to acknowledge the support of the Norwich Bioscience Institutes Research Computing team and the Technical Genomics group at the Earlham Institute, supported by UK Research and Innovation, Core Capability Grant BB/CCG1720/1.

---

**Supplementary Table 1;** Subpopulation, phenotypic annotation for the common bean accessions used in this study.

**Supplementary Table 2;** Genes and non-synonymous genetic variants within each of the identified QTLs associated with determinacy.

**Supplementary Table 3;** Genes and non-synonymous genetic variants within each of the identified QTLs associated with photoperiod sensitivity.

